# IL-17A rescues motor deficits in a mouse model of Spinocerebellar Ataxia Type 2

**DOI:** 10.64898/2026.03.31.715603

**Authors:** Yasmin Yarden-Rabinowitz, Changhyeon Ryu, Ching-Tzu Huang, You-Hyang Song, Yossef Yarom, Gloria B. Choi

## Abstract

Motor performance and coordination deficits are hallmarks of spinocerebellar ataxias, yet effective disease-modifying therapies remain limited. Here, we investigate the expression of the interleukin 17 receptor subunit A (IL-17RA) in cerebellum and assess the therapeutic potential of its ligand in a mouse model for Spinocerebellar Ataxia Type 2 (SCA2). We found that IL-17RA is highly enriched in cerebellar molecular layer interneurons (MLIs), which provide inhibitory input to Purkinje neurons. *In-vitro* electrophysiological recordings revealed that symptomatic SCA2 mice exhibited increased spontaneous inhibitory synaptic input onto Purkinje neurons, which was normalized by IL-17A application to control levels. Concomitantly, IL-17A application restored Purkinje neuron firing, a parameter characteristically reduced in SCA2 mice. Behaviorally, intranasal administration of IL-17A restored motor performance of symptomatic SCA2 mice to control levels in both rotarod and beam-crossing assays. Collectively, our results indicate that IL-17A rescues Purkinje neuron dysfunction and motor deficits in SCA2 mice, highlighting IL-17A signaling as a promising therapeutic target for spinocerebellar ataxia.

## Introduction

Ataxia is a neurodegenerative disorder characterized by impaired movement, coordination, and balance^1^. Among the various forms of ataxia, Spinocerebellar Ataxia Type 2 (SCA2) is a particularly severe inherited neurodegenerative form of the disorder^2,3^, with an estimated global prevalence of 1-2 cases per 100,000 individuals^4^. SCA2 is caused by an abnormal expansion of CAG repeats in the ATXN2 gene, resulting in a polyglutamine (polyQ) expansion that contributes to neuronal dysfunction and degeneration. This neuropathology primarily affects the cerebellum, brainstem, and spinal cord, with cerebellar Purkinje neurons (PNs) exhibiting the most prominent neurodegeneration, characterized by reduced firing rates and premature cell death^3^. Currently, there is no cure for SCA2, and available treatments aim to alleviate symptoms and slow the decline in motor function by stabilizing Purkinje neuron activity^5^. Because Purkinje neuron activity directly regulates cerebellar output and motor control^6–12^, this neuronal dysfunction directly underlies the motor deficits observed in SCA2^13–17^. To investigate SCA2 disease mechanisms and therapeutic approaches, we used transgenic ATXN2^Q127^ mice, a well-established SCA2 model^16^. This model expresses full-length human ATXN2 with 127 CAG repeats specifically in cerebellar Purkinje neurons under control of the Pcp2/L7 promoter, enabling us to examine neuronal activity and motor performance.

The IL-17 family of cytokines, primarily produced by lymphocytes including T and innate lymphoid cells, are immune effector molecules^18^. IL-17A is the predominant cytokine of the IL-17 family (IL-17A–F) and is chiefly produced by Th17 cells, a specialized subset of CD4+ helper T cells. While IL-17A plays a protective role against extracellular bacterial and fungal infections^19^, it is also a pathogenic driver of chronic inflammation, contributing to tissue damage and immune dysregulation across a range of autoimmune and inflammatory diseases^20–22^. However, under physiological conditions, IL-17A serves essential homeostatic functions by coordinating balanced immune responses, including controlled cytokine production, neutrophil recruitment, and antimicrobial defense. This dual nature, from detrimental inflammatory mediator to protective immune regulator, underscores the critical importance of maintaining appropriate IL-17A levels for immune equilibrium^23^. Beyond its established immunological functions, IL-17A has recently been implicated as a neuromodulator capable of influencing neuronal activity in the brain. Specifically, recent work has shown that IL-17A modulates neuronal activity in neurons expressing IL-17RA, leading to behavioral changes such as alterations in social interaction or anxiety^24–27^.

IL-17 receptor subunit a (IL-17RA) is the obligatory subunit required for binding the IL-17A ligand^28^. Our investigation into IL-17A receptor expression in the cerebellum revealed a selective enrichment in cerebellar MLIs. The principal function of the molecular layer is to provide inhibition onto Purkinje neurons, the sole output of the cerebellar cortex. Recordings of spontaneous inhibitory postsynaptic currents (sIPSCs) in Purkinje neurons in ATXN2^Q127^ mice revealed a rightward shift in amplitude distribution relative to controls, indicating that ataxic Purkinje neurons receive stronger inhibitory input, similar to what has been previously reported in another form of SCA^29^. Notably, application of IL-17A shifted the distribution of sIPSC amplitudes toward control values and increased the spontaneous firing rates of ATXN2^Q127^ Purkinje neurons, restoring them to levels comparable to controls. In parallel, we observed that intranasal administration of IL-17A rescued motor deficits in ATXN2^Q127^ mice. Together, our findings highlight a potential immune-based therapeutic strategy for alleviating motor deficits in SCA2.

## Results

### IL-17 receptor subunit a is expressed in the cerebellum

To characterize IL-17RA expression in the cerebellum, we utilized a CRISPR-Cas9-engineered mouse line in which Cre recombinase is expressed under the control of the IL-17RA promoter (IL-17RA-Cre), as previously described^30^. We injected a Cre-dependent EGFP-expressing virus into lobule X in the cerebellar vermis of these mice to visualize IL-17RA-expressing neurons (Figure 1A). EGFP expression was observed in 61% of MLIs, absent from Purkinje neurons (Figure 1B and C) and sparsely in the granule layer (<0.5%, data not shown). We also observed EGFP expression in pinceau^31^, a cone-like synapses surrounding the proximal axonal segment of Purkinje neuron, formed by basket cells, of MLIs (Figure 1B). Approximately 67% of Purkinje neuron somas were surrounded by EGFP-labeled pinceaux at the proximal axon initial segment.

**Figure 1:**
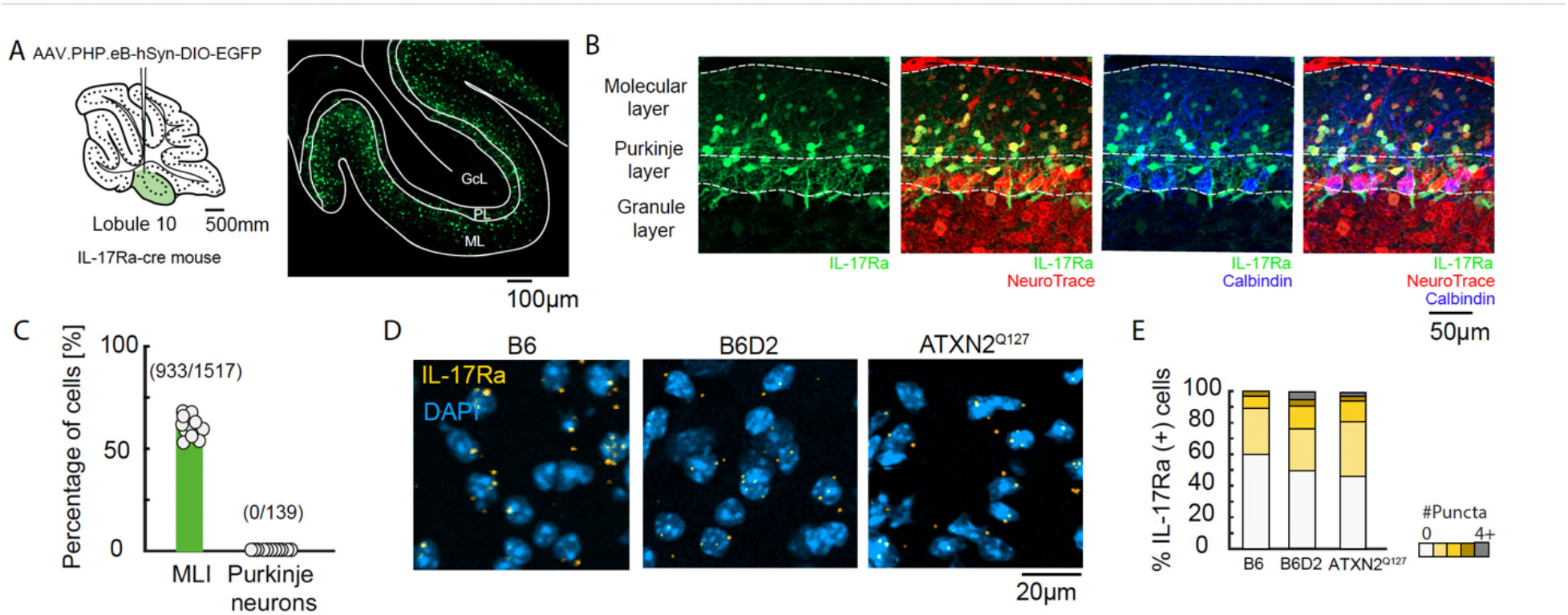
IL-17RA is expressed in molecular layer interneurons. (A) Left: Schematic of the experimental design to drive the expression of EGFP in IL-17RA-expressing cerebellar neurons in a Cre-dependent manner by locally injecting a virus (AAV.PHP.eB-hSyn-DIO-EGFP) into cerebellar lobule X of the IL-17Ra-cre mouse line. Right: Representative EGFP-expressing coronal cerebellar section showing EGFP-labeled neurons (green) within lobule X (ML: Molecular layer, PL: Purkinje layer, GcL: Granule cell layer). (B) Representative EGFP-expressing neurons (green) in a sagittal section of the cerebellar cortex (vermis, lobule X). Inhibitory neurons in the molecular layer are identified with NeuroTrace (red), and Purkinje neurons are labeled with anti-calbindin (blue). Scale bar=50μm. (C) The percentage of EGFP-positive cells relative to the total number of marker-labeled cell type in vermis lobule X Molecular and Purkinje layers: NeuroTrace for molecular layer interneurons, anti-calbindin for Purkinje neurons (n=21 ROIs, 1 ROI per slice, 3 slices per animal, 3 animals; 3 independent experiments) (D) Representative images of *IL-17ra* mRNA expression (yellow) in vermis lobule X of B6, B6D2 and ATXN2^Q127^ mice together with DAPI (blue) in the molecular layer. (E) Quantification of *IL-17ra* mRNA expression was performed by categorizing cells according to the number of *IL-17ra* puncta detected (0, 1, 2, 3, or ≥4). For each category, the percentage of cells was calculated as the number of cells with that puncta count divided by the total number of DAPI-positive cells. The distribution of puncta across mouse lines was not significantly different (χ^2^= 14.28, p = 0.07). (n=9 ROIs from 1 ROI per animal, 9 animals; 9 independent experiments).

We next asked whether IL-17RA is similarly expressed in ATXN2^Q127^ mice compared to wild-type mice. To answer this question, we performed *in situ* hybridization for *IL-17ra* mRNA expression in ATXN2^Q127^ and compared it with both C57BL/6J (B6) mice, serving as the background strain for the IL-17RA-Cre line, and B6D2F1/J (B6D2) mice, serving as the background strain for the ATXN2^Q127^ mice. Our results indicated that *IL-17ra* mRNA expression in the molecular layer is comparable across all three mouse lines, with approximately half (B6: 47%, B6D2: 50%, ATXN2^Q127^: 53%) of the cells in each group containing at least one *IL-17ra* mRNA punctum (Figure 1E–F).

### IL-17A application normalizes inhibitory post-synaptic currents in Purkinje neurons

MLIs form spatially structured networks converging to shape the temporal and spatial patterns of Purkinje cell activity^32^. Interestingly, a recent study reported that the spontaneous inhibitory postsynaptic currents (sIPSCs) in spinocerebellar ataxia type 7 exhibit increased amplitudes compared to controls^29^. Following this, we asked whether a similar phenotype is also present in SCA2 mouse model. We recorded Purkinje neurons intracellularly in acute cerebellar sagittal slices from 10-week-old ATXN2^Q127^ and control mice using whole-cell voltage-clamp (Figure 2A), measuring sIPSCs at 0 mV at least 1 hour following vehicle application to the bath. The cumulative distribution function (CDF) of sIPSC amplitudes and inter-event intervals (IEIs) was calculated (Figure 2D-E). It revealed that the CDF curve of the ATXN2^Q127^-vehicle group exhibits a rightward shift compared to the vehicle-treated control group (Figure 2D; light purple vs. light blue), indicating increased sIPSC amplitudes. Notably, IL-17A treatment shifted the ATXN2^Q127^ CDF curve leftward (purple, Figure 2D), making it comparable to that of the control groups. In contrast, the CDF of sIPSC inter-event intervals was not significantly different between ATXN2^Q127^ and control groups and was not affected by IL-17A treatment (Figure 2E), suggesting that IL-17A does not alter inhibitory synaptic event frequency.

**Figure 2:**
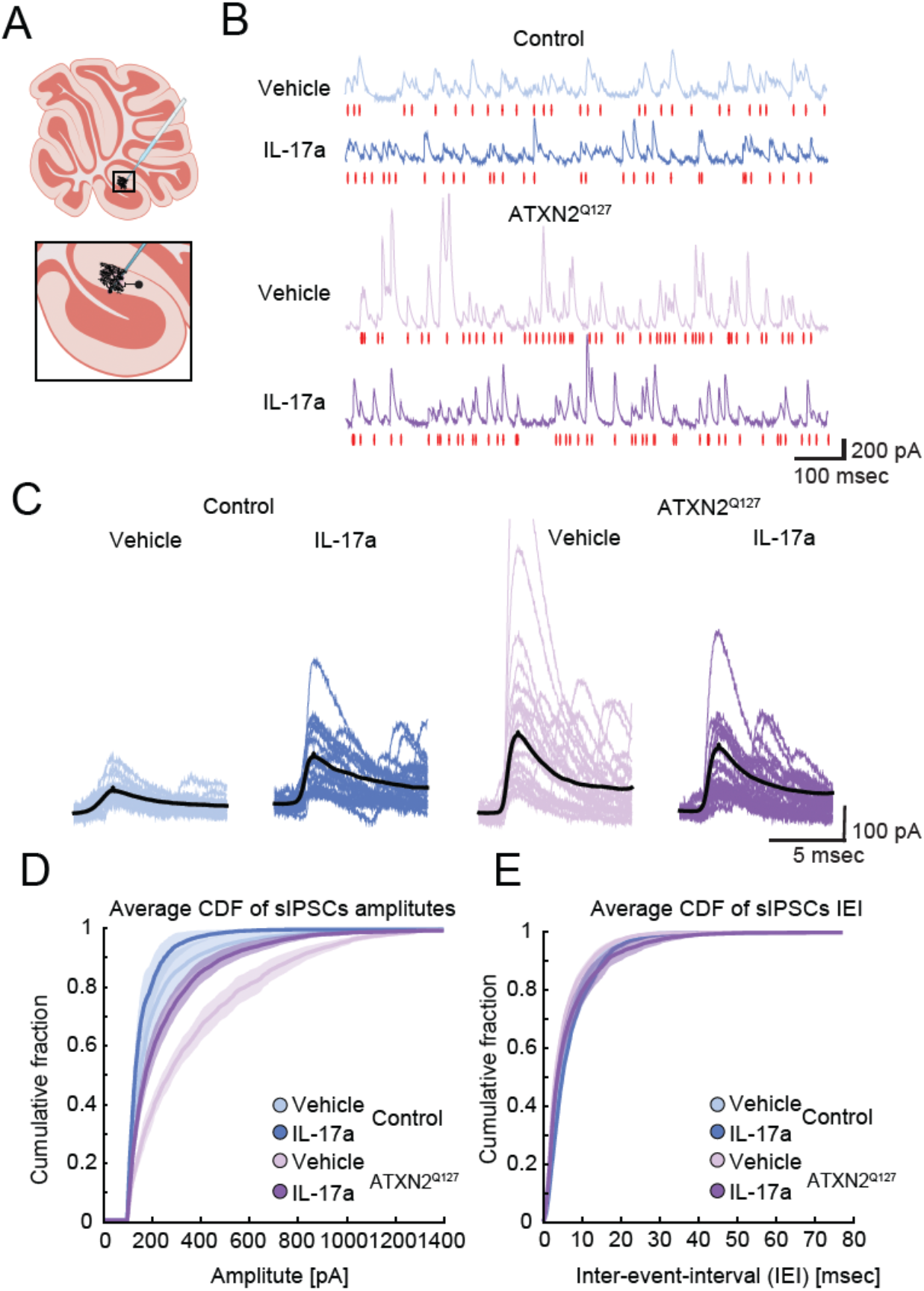
IL-17A decreases inhibitory post-synaptic currents in ATXN2^Q127^ Purkinje neurons. (A) Schematic of intracellular recording of inhibitory synaptic activity in Purkinje neurons in sagittal cerebellar slices from lobule X. (B) Representative current traces of spontaneous inhibitory post-synaptic currents (sIPSC) recorded in Purkinje neurons. Purkinje neurons were recorded in acute cerebellar slices from 10-week-old ATXN2^Q127^ and control mice at least 1 hour following vehicle or IL-17A application to the bath, and whole-cell voltage-clamped sIPSCs were measured at 0 mV. Red lines indicate detected sIPSC events. (C) 20 randomly selected sIPSC events and the mean sIPSC waveform (Black) for each experimental group. (D) The cumulative distribution function (CDF) for sIPSC amplitudes was calculated following vehicle or IL-17A application. (E) CDF for sIPSC inter-event-intervals (n=21 Purkinje neurons recorded from n=2 ATXN2^Q127^ mice, vehicle: n=11, IL-17A: n=10. n=18 Purkinje neurons recorded from n=2 control mice, vehicle: n=9, IL-17A: n=9. n =4 independent experiments).

### IL-17A application rescues Purkinje neuron firing rates and regularity

To understand if the changes in inhibitory input onto Purkinje neurons affect their activity, cell-attached extracellular recordings were obtained from Purkinje neurons in acute cerebellar sagittal slices from 10-week-old ATXN2^Q127^ and control mice (Figure 3A). Our recordings recapitulate previously published results^16^ demonstrating that Purkinje neurons in ATXN2^Q127^ mice fire at lower rates compared to controls. Application of IL-17A increased Purkinje neuron firing rates to values similar to those observed in control neurons (Figure 3B-C).

**Figure 3:**
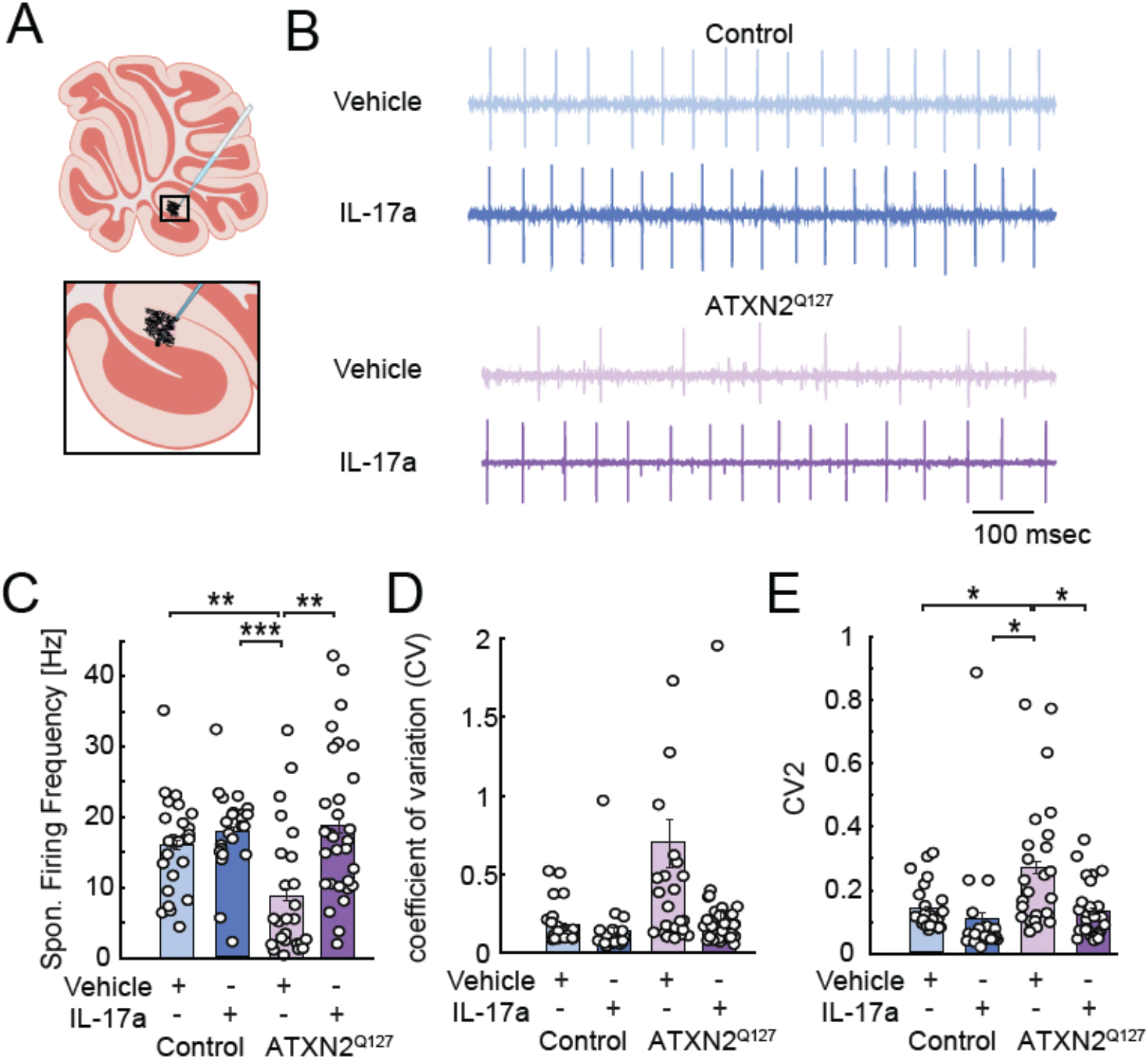
IL-17A increases Purkinje neuron firing rate in ATXN2^Q127^ mice. (A) Schematic of extracellular recording of Purkinje neuron activity in sagittal cerebellar slices from lobule X. (B) Representative traces of spontaneous Purkinje neuron activity 1 hour following vehicle or IL-17A application in control (upper) and ATXN2^Q127^ mice (lower). (C) Mean firing frequency of Purkinje neurons. (D) Coefficient of variation (CV) of interspike intervals for Purkinje neurons. (E) Local Coefficient of Variation (CV2) of interspike intervals for Purkinje neurons (n=55 Purkinje neurons recorded from n=3 ATXN2^Q127^ mice, vehicle: n=25, IL-17A: n=30: n=49 Purkinje neurons recorded from n=2 control mice, vehicle: n=26, IL-17A: n=23. n =5 independent experiments). *p<0.05, **p<0.01, ***p<0.001 as calculated by two-way ANOVA with Sidak’s post-hoc tests.

**Figure 4:**
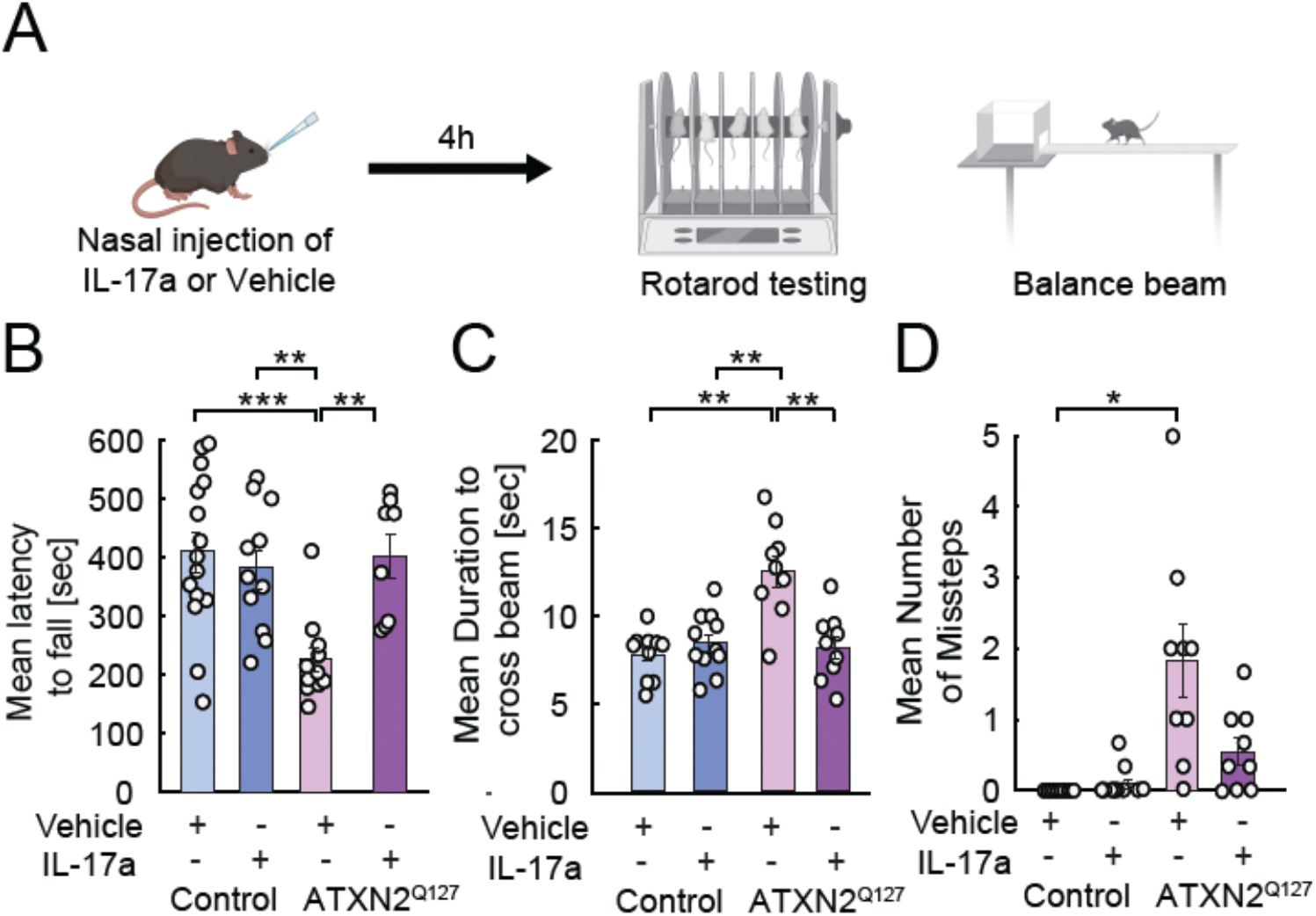
Intranasal administration of IL-17A rescues motor deficits in ataxic mice. (A) ATXN2^Q127^ or control mice were administered vehicle or IL-17A intranasally, and motor performance was tested 4 hours after administration on the accelerating Rotarod and Balance beam tests. (B) Mean latency to fall of the accelerating rotarod 4h after i.n. administration of vehicle or IL-17A in control and ATXN2^Q127^ mice. Individual data points represent mean performance of 3 trials of each mouse (n=19 ATXN2^Q127^ mice, vehicle: n=11, IL-17A: n=8: n=26 control mice, vehicle: n=15, IL-17A: n=11. n =12 independent experiments). (C) Mean duration to cross a 10mm beam 4h after i.n. injection of vehicle or IL-17A in control and ATXN2^Q127^ mice. Each point represents the mean of 3 trials per mouse. (D) Same as in C for the number of missteps (n=18 ATXN2^Q127^ mice, vehicle: n=9, IL-17A: n=9: n=21 control mice, vehicle: n=10, IL-17A: n=11. n =12 independent experiments). *p<0.05, **p<0.01, ***p<0.001 as calculated by two-way ANOVA with Sidak’s post-hoc tests.

We also asked whether IL-17A application affects spiking regularity, as MLIs are known to modulate Purkinje neuron firing patterns^31,32^. To do so, we calculated the coefficient of variation (CV) of inter-spike interval (ISI) and the local coefficient of variation (CV2). CV2 is computed between pairs of consecutive ISIs, making it more sensitive to local irregularities in firing than global variability, which is better captured by CV. Our results indicate that CV2, which is elevated in ATXN2^Q127^ mice under control conditions, is reduced following IL-17A treatment, suggesting that Purkinje neuron firing becomes more regular.

### Intranasal administration of IL-17A to the brain rescues motor deficits in ataxic mice

Previous work established that correcting Purkinje neuron firing improves motor deficits in ataxia^5,33^. Therefore, considering the normalization of firing rates and regularity, we asked whether application of IL-17A could correct motor deficits in ataxic mice. To answer this question, we delivered IL-17A intranasally (i.n.)^27^ and tested motor performance 4 hours after i.n. administration. We used two cerebellar-dependent motor performance tests (Figure 3A). Our results replicate previously published findings that 10-week-old ATXN2^Q127^ mice exhibit motor deficits, falling off the rotarod earlier than controls^16^ and taking longer to traverse the balance beam with increased missteps, consistent with impaired balance and coordination. Notably, i.n. administration of IL-17A rescued these deficits; treated ATXN2^Q127^ mice remained on the rotarod for longer durations (Figure 3B) and crossed the balance beam more quickly (Figures 3C). Across both assays, IL-17A restored motor performance in ATXN2^Q127^ mice to levels comparable to those of control animals.

## Materials and methods

### Animals

All experiments were performed according to the Guide for the Care and Use of Laboratory Animals and were approved by the National Institutes of Health and the Committee on Animal Care at Massachusetts Institute of Technology. C57BL/6 and B6D2F1/J mice were purchased from the Jackson Laboratories and inbred in house. IL-17RA-Cre line was generated using the CRISPR-Cas9 system (Biocytogen). ATXN2^Q127^ mice were obtained as a generous gift from Dr. Stefan M. Pulst (University of Utah, Salt Lake City, UT, USA) and maintained by breeding each generation carrying the transgene with non-transgenic B6D2F1/J mice, following a breeding scheme similar to that described previously^34^. All mice used in this study were males aged 10-14 weeks.

### Stereotaxic surgery

All surgeries were performed under aseptic conditions. Mice were anesthetized with isoflurane and received a preoperative injection of slow-release buprenorphine (1 mg/kg, subcutaneous) at the start of surgery. To visualize IL-17RA-positive neurons, a craniotomy was performed at the following stereotaxic coordinates: 0 mm ML, –6.7 mm AP, and 2.5 mm DV. An AAV.PHP.eB-hSyn-DIO-EGFP virus (Addgene #50457-AAV2) was injected into IL-17RA-Cre mice at the specified coordinates using a pulled fine-glass capillary at a rate of 0.1 μl/min. Following surgery, the virus was allowed to incubate for 2–3 weeks prior to tissue collection or analysis.

### Immunohistochemistry

Animals were transcardially perfused with cold PFA (4% in PBS). Brains were kept in 4% PFA overnight at 4°C prior to vibratome sectioning. Brains were sectioned sagittally at 100 μm-thick. Sections were permeabilized with blocking solution (0.4% Triton X-100 and 2% goat serum in PBS) for 1 h at room temperature. Sections were incubated overnight in a blocking solution containing primary antibodies at 4°C. Primary antibodies used were NeuroTrace (1:1000, ThermoFisher) for molecular layer interneurons, and anti-Calbindin (1:500, Swant) for Purkinje neurons. The following day, sections were incubated with corresponding secondary antibodies, anti-mouse Alexa 647 and\or anti-rabbit Alexa 568 (1:250, ThermoFisher) for 2 hrs at room temperature with Neurotrace (1:500, Invitrogen) and mounted in CC/Mount (Sigma) mounting medium. Images of stained slices were acquired using a confocal (LSM900, Carl Zeiss) with a 20x objective lens. Total number of cells expressing the primary antibody target and double-positive cells expressing both the primary antibody target and EGFP, indication a IL-17RA-cre expressing cell transfected by virus, were detected using QuPath^35^.

### In-situ hybridization

Mice were euthanized by cervical dislocation. Brains were extracted, embedded in optimal cutting temperature (OCT) compound, and placed in 2-methylbutane (M0167, TCI) on dry ice slurry with 100% EtOH in a Styrofoam box. Sections were cut at 10-μm thickness on a cryostat. *In-situ* hybridizations were performed using the RNAscope Multiplex Fluorescent Reagent kit v2 (323100, Advanced Cell Diagnostics) using a probe targeting the IL-17RA (566131, Advanced Cell Diagnostics) as previously described^24^. Sections were counterstained with DAPI. Images were acquired using a confocal microscope (LSM900, Carl Zeiss) with a 10X objective lens. DAPI and IL-17RA puncta expression were quantified using QuPath^35^. Cells were divided into 5 categories based on expression level, 0-3 puncta and 4 or more puncta.

### Acute slice preparations

On the day of the experiment, mice were anesthetized with isoflurane and transcardially perfused with an oxygenated solution containing (in mM): 126 NaCl, 4 KCl, 1 MgCl_2_, 1 CaCl_2_, 1 NaH_2_PO_4_, 26 NaHCO_3_, and 20 glucose, bubbled with 95% O_2_ and 5% CO_2_ at 34–36 °C. The cerebellum was extracted, and 300 μm-thick sagittal sections were sliced using a vibratome (Leica VT1200S) in the same solution and at the same temperature^36^. Slices were then transferred to a chamber containing the same solution maintained at 34 °C for a 60-minute recovery period, after which they were kept at room temperature. All experiments were conducted within 8 hours from the start of the recovery period

### Whole-cell recordings

For recording Purkinje neuron following cytokine or vehicle application, IL-17A (50 ng/ml) or PBS were added to the recording solutions 1 hour prior to recording. Purkinje neurons were patch-clamped using glass pipettes. For recording synaptic activity pipettes were filled with a cesium-based internal solution containing (in mM): 120 Cs-methanesulfonate, 8 NaCl, 10 HEPES, 0.5 EGTA, 10 Na-phosphocreatine, 4 Mg-ATP, 0.4 Na2_22-GTP, 1 QX-314 bromide, and 10 sucrose. The solution was adjusted to pH 7.25 with CsOH and had an osmolarity of ∼294 mOsm. The membrane properties were measured in response to hyperpolarizing current pulses (100 ms duration, -5 mV) in the voltage-clamp mode. For extracellular cell-attached recordings of Purkinje neurons, we used a solution containing (in mM): 125 NaCl, 3 KCl, 2 CaCl_2_ and 10 HEPES. The solution was adjusted to pH 7.25 with NaOH and had a final osmolarity of ∼290–295 mOsm. Voltage-and current-clamp recordings were performed in whole-cell configuration using a patch-clamp amplifier (Molecular Devices) and a Digidata 1550B digitizer (Molecular Devices) controlled by pCLAMP software (Molecular Devices).

### Exclusion inclusion criteria

Recording of IPSCs: Cells were held at 0 mV; recordings that required holding currents more negative than 400 pA were excluded. Access and membrane properties were monitored with 5-mV test pulses; recordings showing capacitive transients <300 pA or test-pulse response amplitudes >100 pA were rejected.

### Identifying sIPSCs

Spontaneous IPSCs were detected and analyzed offline using MATLAB. Event amplitudes were calculated as the difference between the preceding local minimum and the subsequent local maximum of each detected event. Only events with amplitudes greater than 100 pA were included in the analysis. Recordings segments were considered stable and included only if the mean event amplitude and inter-event-interval during the first and last 20% of the analyzed segment did not differ by more than 20%.

### Calculating CV2

To quantify local variability in Purkinje neuron firing, we calculated CV2 for each pair of consecutive inter-spike intervals (ISIs). CV2 was computed as:

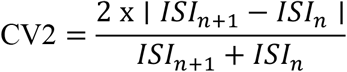

### Rotarod Test

Motor coordination and balance were assessed using the accelerating rotarod test (Ugo Basile, Italy). Mice were placed on a rotating rod with an initial speed of 4 rpm, which gradually accelerated to 40 rpm over a 10-minute period. Each animal underwent three trials with an intertrial interval of at least 20 minutes. The latency to fall was recorded for each trial, and the average performance was used for analysis. Mice that passively clung to the rod and completed 3 full rotations without active locomotion were scored as having fallen at the time of the first passive rotation.

### Balance Beam Test

The apparatus consisted of a narrow wooden beam (100 cm in length, 1 cm in width) elevated 20 cm above the surface. On the day of the experiment, mice were first habituated to the goal box by placing it in their home cage for 20 minutes. Subsequently, animals were acclimated to the apparatus once by being guided to walk the final third of the balance beam, enter the goal box, and remain there for 30 seconds. Acclimation was performed prior to the first test trial. During testing, the latency to traverse the beam was recorded using a photogate (Eisco Scientific; Catalog No. 470317-428) and the number of hindlimb slips was recorded manually by the experimenter. A second experimenter independently repeated the experiments to assess and confirm the reproducibility of the measurements.

## Discussion

In this study, we show that application of the cytokine IL-17A to a mouse model of SCA2 modulates inhibitory input onto Purkinje neurons, restoring both firing rates and regularity. Consistent with prior work showing that normalization of Purkinje neuron activity improves motor deficits in ataxia^33^, our findings further show that IL-17A treatment alleviates motor deficits in SCA2 mice.

At the circuit level, Purkinje neuron firing rates were significantly reduced in SCA2 mice, consistent with previous reports^37^. And we observed a rightward shift in the cumulative distribution of Purkinje neurons’ sIPSC amplitudes in these mice, indicating enhanced inhibitory drive. Together, these observations suggest that increased inhibitory inputs may contribute to the decrease in Purkinje neuron firing.

Beyond firing rate, the regularity of Purkinje neuron firing has been proposed to play a critical role in cerebellar information processing by supporting population-level spatiotemporal activity patterns that complement rate-based coding^38,39^. Subsequent studies have directly linked Purkinje neuron firing regularity to movement kinematics^40^, and these regular firing patterns are thought to be shaped by molecular layer inhibitory interneurons^32^. Consistent with this framework, we found that CV2, a measure of variability between successive interspike intervals, was elevated in SCA2 mice, indicating increased firing irregularity. These observations align with prior findings in SCA2^41^ and ataxia more broadly^33^. Notably, IL-17A application reduced this irregularity in Purkinje neurons. In this regard, while the precise mechanism of IL-17A remains to be defined, especially with respect to downstream signaling within receptor-expressing neurons such as molecular layer interneurons, previous studies have likewise demonstrated that IL-17A reduces inhibitory input^42^ and have suggested distinct mechanisms through which synaptic transmission may be more broadly affected^43,44^.

In summary, our results demonstrate that IL-17A restores Purkinje neuron firing dynamics and improves motor performance in SCA2 mice. These findings support the idea that modulation of cerebellar circuitry via IL-17A signaling represents a promising therapeutic strategy for spinocerebellar ataxias. More broadly, this work suggest that immune-derived signals can exert functionally significant effects on cerebellar circuits, with potential relevance for other neurodegenerative disorders characterized by Purkinje neuron dysfunction.

### Limitations of the study

Our work demonstrates robust IL-17RA expression in molecular layer interneurons, and that the inhibitory input onto their postsynaptic cells is modulated following IL-17A application. However, in the scope of this work, we did not functionally manipulate MLIs, and future studies employing cell-type-specific conditional knockouts of IL-17RA in MLIs will be necessary to delineate their precise contribution. Moreover, under *in-vivo* conditions, we cannot exclude the contribution of other IL-17RA-expressing cell types in interconnected brain regions, which may also play a role in the presented IL-17A rescue of motor deficits.

